# Phase separation of the LINE-1 ORF1 protein is mediated by the N-terminus and coiled-coil domain

**DOI:** 10.1101/2020.10.29.360719

**Authors:** JC Newton, GY Li, MT Naik, NL Fawzi, JM Sedivy, G Jogl

## Abstract

Long Interspersed Nuclear Element-1 (LINE-1 or L1) is a retrotransposable element that autonomously replicates in the human genome, resulting in DNA damage and genomic instability. Activation of L1 in senescent cells triggers a type I interferon response and age-associated inflammation. Two open reading frames encode an ORF1 protein functioning as mRNA chaperone and an ORF2 protein providing catalytic activities necessary for retrotransposition. No function has been identified for the conserved, disordered N-terminal region of ORF1. Using microscopy and NMR spectroscopy, we demonstrate that ORF1 forms liquid droplets *in vitro* in a salt-dependent manner and that interactions between its N-terminal region and coiled-coil domain are necessary for phase separation. Mutations disrupting blocks of charged residues within the N-terminus impair phase separation while some mutations within the coiled-coil domain enhance phase separation. Demixing of the L1 particle from the cytosol may provide a mechanism to protect the L1 transcript from degradation.

**Statement of significance:** Over half of the human genome is comprised of repetitive sequences. The Long Interspersed Nuclear Element-1 (L1) is an autonomous mobile DNA element that can alter its genomic location, resulting in genomic instability and DNA damage. L1 encodes two proteins that are required for this function: the ORF1 RNA chaperone and the enzymatic ORF2. Here, we demonstrate that ORF1 forms liquid-liquid phase separated states *in vitro*, which is mediated by electrostatic interactions between the conserved, disordered N-terminus and coiled-coil domain. This work provides a framework to explore how L1 phase separation may enhance the ability of the retrotransposable element to colonize the genome by preventing degradation of the L1 transcript and evasion of host immune responses.

## Introduction

Over half of the human genome is composed of repetitive sequences (1), the largest fraction of which are mobile DNA sequences known as retrotransposable elements (RTEs) (2). They are comprised of three major classes, the Long Terminal Repeat (LTR) elements (related to retroviruses) and the non-LTR elements, known as Long Interspersed Nuclear Elements (LINEs) and Short Interspersed Nuclear Elements (SINEs). RTEs colonize their host genomes using a ‘copy-and-paste’ mechanism that employs an RNA intermediate and an RTE-encoded reverse transcriptase enzyme (3). RTEs can exert deleterious effects on their host cells in many ways, including insertional mutagenesis, promotion of DNA damage and genomic structural instability, triggering of epigenetic changes, and disruption of normal patterns of gene regulation (4). The only autonomously active human RTEs are the LINE-1 elements (L1). L1s comprise approximately 17% of the human genome (~500,000 copies), but only the most evolutionarily recent subfamily L1HS has active members (5, 6). Until recently, L1s were thought to be silent in the soma, but new evidence points to activity in the brain (7), cancers (8), cellular senescence (9) and aging (10–12). In senescent cells, transcriptional activation of L1 and the ensuing reverse transcription of its cDNA triggers a type I interferon response, which promotes age-associated inflammation, also known as inflammaging (11, 12). Given that inflammaging has been linked with several important age-related, chronic diseases (13), further understanding of the details of the L1 lifecycle in somatic cells is expected to provide novel therapeutic targets.

L1 is a 6 kb element whose 5’ UTR contains an internal promoter from which two proteins required for retrotransposition are expressed. The 40 kDa ORF1 protein functions as an mRNA chaperone and packing protein while the 150 kDa ORF2 protein encodes the endonuclease (EN) and reverse transcriptase (RT) activities necessary for retrotransposition (Fig. 1A) (3, 14). The L1 ribonucleoprotein (RNP) particle assembles in *cis* in the cytoplasm before entering the nucleus for integration into a novel site in the host genome (15). ORF2 binds the L1 transcript at its 3’ end while the ORF1 protein coats the remainder of the mRNA (16). Retrotransposition in the nucleus occurs by a mechanism known as target-primed reverse transcription (TPRT) which is initiated by an EN-generated nick in the genome (2). The displaced flap of DNA binds to the poly A tail of the mRNA template and is then reverse transcribed at the site of insertion. To enable this life cycle, it is necessary to maintain an intact, full-length transcript for successful TPRT, which highlights the importance of the ORF1 chaperone functions (14, 17).

**Figure 1.**
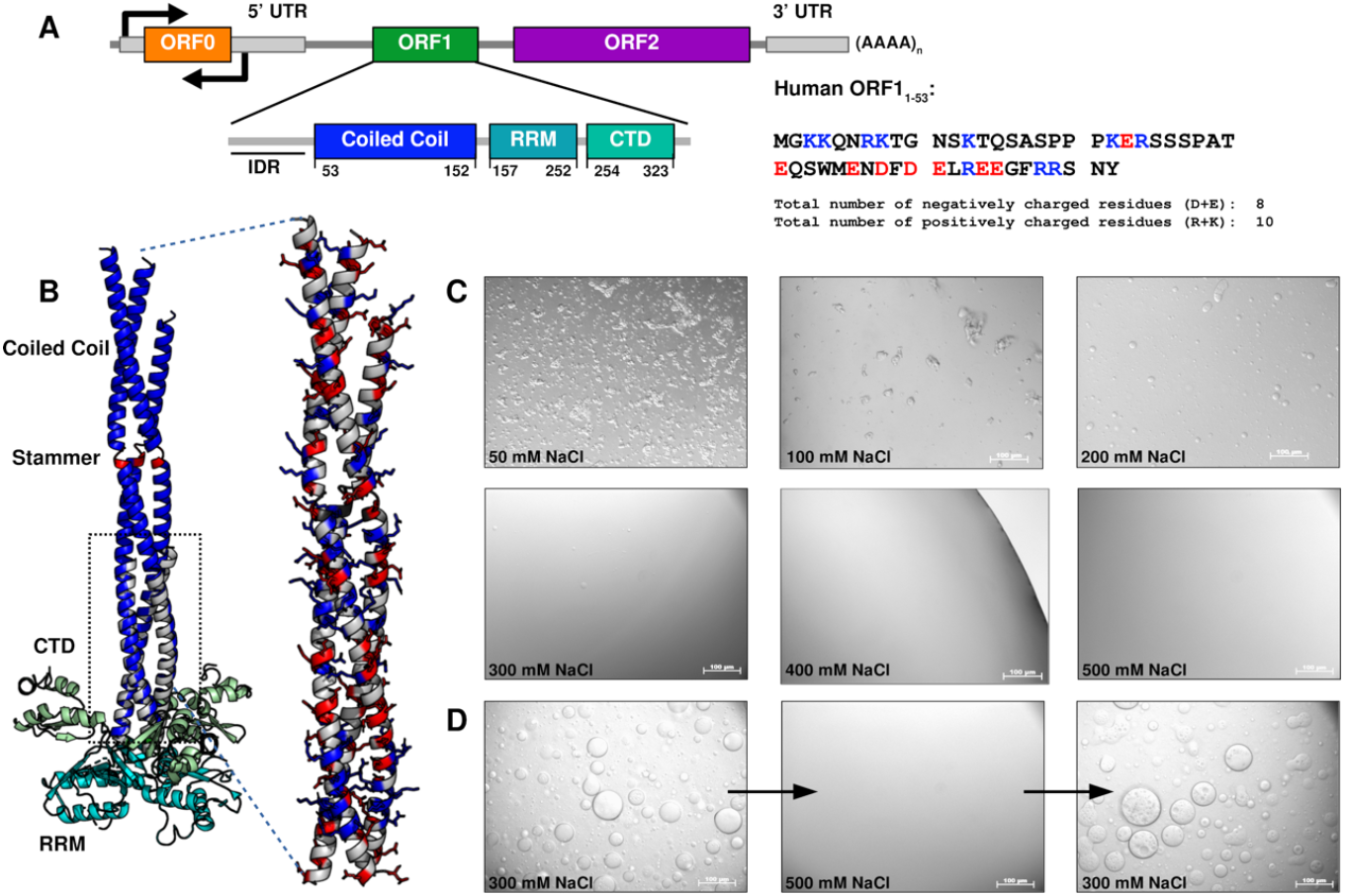
ORF1 phase separation is driven by electrostatics and is reversible. **A.** Domain organization of L1 and ORF1 functional units. ORF1 consists of three folded domains: a coiled-coil region responsible for trimerization of the ORF1 protein, an RNA recognition motif (RRM), and a C-terminal domain (CTD), that act in concert to bind to L1 RNA. The intrinsically disordered N-terminal region is denoted as IDR. The sequence of the N-terminal 53 residues is shown color-coded by charge. **B.** Composite model of the ORF1 trimer. The extended coiled-coil domain (PDB entry 6FIA) is superposed on the ORF1 core structure (PDB entry 2YKO) in the boxed region. Domains are colored as shown in A. The stammer region is highlighted in red. An enlarged view of the coiled-coil domain shows the charge distribution on the solvent exposed surface with basic residues shown in blue and acidic residues in red. **C.** The extent and morphology of the ORF1 phase-separated state is dependent on the NaCl concentration in the buffer solution. In low salt concentrations, ORF1 forms amorphous aggregates while forming spherical droplets in ranges of 200-400 mM NaCl. In 500 mM NaCl or higher, ORF1 remains dispersed. **D.** ORF1 phase separation is reversible. Increasing the NaCl concentration from 300 mM to 500 mM while maintaining protein concentration (300 μM) solubilizes phase separated ORF1. Phase-separated droplets reappear when the NaCl concentration is reduced to 300 mM.

Two crystal structures of the ORF1 protein have been determined and indicate that ORF1 trimerizes via its coiled-coil domain (18, 19) (Fig. 1B). The C-terminal portion of the protein contains an RNA Recognition Motif (RRM) and the C-terminal domain (CTD). Together, these two domains form a positively charged surface that is predicted to weave the single-stranded L1 RNA around the base of the ORF1 trimer (18, 19).

The 150 Å long coiled-coil domain is organized in a series of 14 heptads with small hydrophobic residues in positions two and four that orient to the center of the helical bundle formed by the trimer. The C-terminus of the coiled coil contains seven heptads with repeating R-h-x-x-h-E motifs (where x denotes any amino acid and h denotes the hydrophobic residues at positions 2 and 4) that form salt bridges to stabilize trimerization. This sequence is highly conserved across species (18). Residues 91-93 (MEL) are inserted within heptad 9 and form a three-residue bulge, or stammer, that destabilizes the N-terminal portion of the coil and contributes to an observed metastability that seems to be required for L1 retrotransposition (19) (Fig. 1B). This suggests that the N-terminal portion of the coil can switch between fully structured or partially unstructured states, which is corroborated by lacking electron density for heptads 13-14 within the shortest helix in the crystal structure (19). Despite the fact that ORF1 has been well characterized structurally, less is known about the function of the N-terminal region. Mutations in ORF1 that disrupt nucleic acid binding, deletion of the stammer insertion in the coiled-coil domain, or removal of the positive charge at the N-terminus all drastically impair retrotransposition, to an extent similar to abrogating RT activity (such as the D702Y mutation (18–21)). In addition, expression of truncated ORF1 proteins containing the N-terminal and coiled-coil region suppress L1 retrotransposition, supporting the importance of these domains for L1 function (22).

In cell-culture overexpression systems, the ORF1 protein has been observed within two membrane-less organelles, stress granules and processing bodies (P-bodies), in the cytoplasm of cells both in the presence and absence of cellular stress (23, 24). Tandem affinity capture of fully assembled cytoplasmic L1 RNP particles, using a FLAG antibody against tagged ORF2 followed by a monoclonal ORF1 antibody, identified interactions between several stress granule or P-body associated proteins and the fully assembled cytoplasmic L1 RNP (25–28). It is possible that stress granules provide an additional layer of protection from circulating proteases, RNases, and immune sensors and give the L1 RNP complex additional time to properly form. This phenomenon has been observed for the assembly of the LTR retrotransposon Ty3 into a capsid-like structure in yeast P-bodies (29). On the other hand, sequestration within stress granules may target the L1 RNP assembly for degradation as an additional host defense response against retrotransposition (23).

The formation stress granules and other membrane-less organelles can be driven by liquid-liquid phase separation (LLPS), often initiated by RNPs, to partition the cellular space and exert spatiotemporal control over biochemical reactions within each particle (30, 31). Weak intermolecular interactions between proteins and nucleic acids drive coalescence into a phase-separated state where biomolecules coexist in condensed and dilute phases. In the condensed phase, these biomolecules are at high local concentrations that function to exclude solvent and other biomolecules in favor of interactions with specific proteins or nucleic acids (32). This forms a chemically distinct environment that is able to reversibly demix from the surrounding cytoplasm or nucleoplasm (33). Because liquid phase-separated states internally consist of highly dynamic interactions and lack a physical membrane barrier, some of these structures are able to rapidly form or disassemble in response to the cellular environment, exemplified by stress granule formation and disassembly during times of heightened cellular stress and subsequent removal of the stressor (32). The ability of proteins to incorporate into coacervates is often encoded in modular domains that enable multivalent interactions or in intrinsically disordered regions that form complexes through intermolecular contacts in a sequence-dependent manner (30, 34). For example, several proteins known to localize to physiological granules and that readily undergo phase separation *in vitro*, such as Fused in Sarcoma (FUS) (35) or TAR DNA-binding protein-43 (TDP-43) (36), have low complexity sequences, which are regions composed of short repeat sequences that mediate non-specific interactions with other biomolecules within the phase-separated body. Furthermore, folded RNA-binding domains as well as disordered segments with arginine-rich sequences are features of both these proteins, contributing to multivalency of interaction (36). While the protein does not contain easily identifiable repeat sequences, many of the critical features of phase-separating proteins are present in ORF1.

Because of the association of ORF1 with cytoplasmic phase-separated bodies, the requirement of its putatively disordered N-terminal domain for L1 retrotransposition, and the multivalent RNA-binding architecture similar to known phase-separating proteins, we hypothesized that the disordered N-terminal region promotes phase separation of the ORF1 protein. Using biophysical phase separation assays and NMR spectroscopy, we show that, while the presence of the N-terminus is necessary for ORF1 condensation, *in vitro* liquid-liquid phase separation is also mediated by the interaction of the disordered N-terminus with the C-terminal portion of the trimeric coiled-coil domain. Interactions that drive ORF1 phase separation are dependent on electrostatics and can be modulated by the solution conditions and by mutations within the ORF1 protein.

## Materials and Methods

### Cloning and expression of recombinant proteins

A plasmid containing the bacterial codon optimized sequence of ORF1_1-338_ was synthesized by GenScript in the pUC57 cloning vector (Supporting Material). ORF1 sequences of interest were PCR amplified using either Thermo Scientific Phusion Hot Start II High-Fidelity PCR Master Mix or ACTGene ACTaq High Fidelity DNA Polymerase. The inserts were cloned into the pET26b vector for expression (Novagen) using NdeI and XhoI double digested products for T4 ligation or undigested insert for Gibson assembly into the pET26b expression vector (37). ORF1 mutants were generated using the pET26b-ORF1_1-338_ as a template for overlap extension PCR as described previously (38). The ORF1 insert was PCR amplified with primers containing the mutation of interest and T7 promoter or terminator primers. The PCR product was gel extracted and the forward and reverse products were mixed as a template for the next round of PCR amplification using the universal T7 promoter and terminator primers. The PCR product was gel extracted and cloned into an empty pET26b vector as described above.

ORF1 expression plasmids were transformed into *Escherichia coli* BL21STAR(DE3) cells (Invitrogen). One-liter cultures supplemented with 50 μg/mL kanamycin were inoculated with 20 mL of an over-night culture and grown at 37°C to an OD_600_ between 0.5 and 0.8. Cultures were cooled for 1 hour to 20°C and 0.5 mM IPTG was added to induce protein expression for 16-18 hours. Cells were harvested in a Beckman SLC-6000 rotor at 5,500 x g for 12 minutes at 4°C.

### ORF1 purification

Cells were resuspended in 10 mL of lysis buffer (20 mM Tris pH 8.0, 1 M NaCl, and 5 mM Imidazole supplemented with PMSF and Benzamidine) per 1 g of dry cell pellet mass and lysed using an Avestin EmulsiFlex C3. Lysate was clarified at 30,000 rpm for 60 minutes at 4°C in a Beckman Ti45 ultracentrifuge rotor and filtered through a 0.22 μm PES membrane before being loaded onto a GE XK 16/20 column packed with 25 mL of Qiagen Ni-NTA resin. The column was washed with a minimum of 5 column volumes of HisTag Buffer A (20 mM Tris pH 8.0, 1 M NaCl, 5 mM Imidazole) and eluted with a gradient of 0-70% HisTag Buffer B (20 mM Tris pH 8.0, 1 M NaCl, 500 mM Imidazole). Fractions corresponding to the ORF1 protein were pooled and concentrated to 5 mL using the Millipore 10 kDa Amicon Ultra Centrifugal Filters and injected onto a GE HiPrep 16/60 Sephacryl S-300 HR gel filtration column equilibrated in 20 mM Tris pH 8.0, 1 M NaCl, 1 mM DTT. All constructs and mutants of ORF1 were purified using the same protocol except ORF1_1-53_, which was heat purified before gel filtration. Once concentrated to 5 mL, ORF1_1-53_ was incubated at 70°C for 15 minutes. The protein was cooled on ice for 15 minutes and any precipitation was removed by centrifugation at 12,000 rpm for 10 minutes at room temperature.

### Microscopy

Samples of ORF1 for differential contrast (DIC) microscopy were prepared by diluting a 1-2 mM concentrated stock of the ORF1 protein into a lower salt buffer. 5 μL of the sample were spotted onto a glass cover slip for imaging using 20X magnification and Nomarski optics on a Zeiss Axiovert 200 M microscope. All microscopy images shown are of 300 μM ORF1 protein in 20 mM Tris pH 8.0, 300 mM NaCl, 1 mM DTT, unless otherwise noted.

For the titration experiments between ORF1_1-53_ and either the full-length ORF1_1-338_ or coiled-coil domain ORF1_53-152_, either 2 mM (1:1), 6 mM (1:3), or 12 mM (1:6) ORF1_1-53_ was added to 2 mM ORF1_53-152_ or ORF1_1-338_ in 20 mM Tris pH 8.0, 1 M NaCl, 1 mM DTT. All samples remained clear upon mixing. They were subsequently diluted to 300 μM, 150 μM, or 50 μM ORF1_53-152_ or ORF1_1-338_ in 500 mM, 300 mM, and 150 mM NaCl buffers. Phase separation was assayed by microscopy as described above.

### Dilute phase diagram

ORF1 was diluted to 300 μM in a final buffer of 20 mM Tris pH 8.0 and 1 mM DTT with salt concentrations of either 100 mM, 150 mM, 200 mM, 250 mM, 300 mM, 350 mM, or 400 mM NaCl. 100 μL samples were incubated at 25 °C in a thermocycler for 30 minutes. The condensed phase was separated from the dilute phase by centrifugation at 10,000 rpm for 30 seconds. Protein concentrations in samples of the dilute phase were measured directly on a Thermo Nanodrop 2000 Spectrophotometer using the appropriate buffer as a blank. Concentrations were calculated using the Beer-Lambert law with the predicted extinction coefficients and molecular weights corresponding to each construct. Errors bars note the standard deviation of three technical replicates.

### NMR Spectroscopy

All NMR experiments were carried out at 25 °C on Bruker Avance III HD 850 MHz or NEO 600 MHz spectrometers equipped with 5 mm TCI cryoprobes and single-axis pulsed-field gradients. NMR data were acquired on isotope enriched ORF1_1-53_ samples (20 μM to 1.6 mM) dissolved either in 20 mM HEPES; pH 7.25 or 20mM MES; pH 6.0 buffers containing 200 mM NaCl, 1 mM DTT and 5% D2O using standard pulse-programs provided by the instrument manufacturer. NMR datasets were acquired and processed using Topspin software, version 3.5 or 4.0 (Bruker, Germany), and were further analyzed in Sparky, version 3.112 (University of California, San Francisco). Backbone resonance assignments for ORF1_1-53_ were achieved through the analysis of standard 3D-triple resonance experiments HNCA, HNCACB, and HNCO. ^15^N relaxation experiments were performed on ^15^N-labeled ORF1_1-53_ or in the presence of equimolar complex with non-labeled ORF1_53-152_. The values of the ^15^N longitudinal relaxation rate constants (*R*_1_) were measured using interleaved experiments with relaxation delays of 20, 40, 60, 100, 200, 400 and 800 ms. The values of the transverse relaxation rate constants (*R*_2_) were measured using interleaved experiments with relaxation delays of 15.7, 31.4, 78.5, 110, 157, 173 and 251 ms. Both rate constants were calculated assuming a monoexponential decay of the peak intensities using the program Curvefit (Prof. Arthur Palmer, Columbia University). The steady-state heteronuclear {^1^H}-^15^N NOE experiment was carried out in an interleaved manner, with and without proton saturation and the NOE effect was calculated as a ratio of the peak intensities.

## Results

### The full-length ORF1 protein phase separates, and this property is dependent on electrostatic interactions

During purification of the recombinant full-length ORF1_1-338_ protein, we observed that protein solubility is strongly dependent on salt concentration and that protein purification requires high salt concentrations to prevent oligomerization. In gel filtration experiments of soluble ORF1_1-338_ in 1 M NaCl buffer, the protein elutes at a position consistent with trimer formation via the coiled-coil domain, as previously described in structural studies of both an RNA-binding construct ORF1_110-321_ and an ORF1_53-152_ construct that isolates the extended coiled-coil domain (18, 19). Reduction of NaCl concentration promotes liquid-liquid phase separation of full-length ORF1_1-338_, and the extent of ORF1 condensate formation is dependent on both NaCl and protein concentrations (Fig. 1C). In buffers with 300 mM NaCl, ORF1 condensates form spherical droplets that behave as dynamic liquids, capable of fusing with neighboring droplets (Fig. S1 in the Supporting Material).

By increasing the NaCl concentration to 500 mM, while maintaining a 300 μM ORF1 protein concentration, ORF1 condensates are dispersed into a single phase with the protein evenly distributed in solution. Reducing the NaCl concentration back to 300 mM in the same sample subsequently reinitiates ORF1 condensation, demonstrating reversibility (Fig. 1D). However, reducing the salt concentration to 50-150 mM NaCl causes the ORF1 protein to form irregularly shaped structures that do not fuse and flow, indicating that the ORF1 phase-separated state is capable of forming either higher order structures or hydrogels. At NaCl concentrations between 500 mM and 1 M NaCl, ORF1 remains well mixed with the surrounding buffer. The observed salt dependency suggests that electrostatic interactions likely play an important role in biomolecular condensation of ORF1_1-338_.

### Truncated ORF1_1-152_, containing the N-terminus and coiled-coil domain, phase-separates readily, with similar characteristics of full-length ORF1 LLPS

In order to identify the domains involved in ORF1 phase separation, we designed constructs that isolated either individual or tandem domains. We found that neither the RRM nor CTD are capable of phase separating on their own or in tandem (data not shown), whereas truncation of the N-terminal 53 or 65 residues largely reduces the extent of phase separation. In contrast, a minimal construct containing the N-terminus and coiled-coil domain (ORF1_1-152_) phase separates readily, similar to the full-length protein and demonstrates the same characteristics of phase separation, such as droplet numbers and size, at lower salt concentration (Fig. 2A, Table 1). Therefore, we chose to analyze molecular interactions driving ORF1_1-152_ phase separation in greater detail.

**Figure 2.**
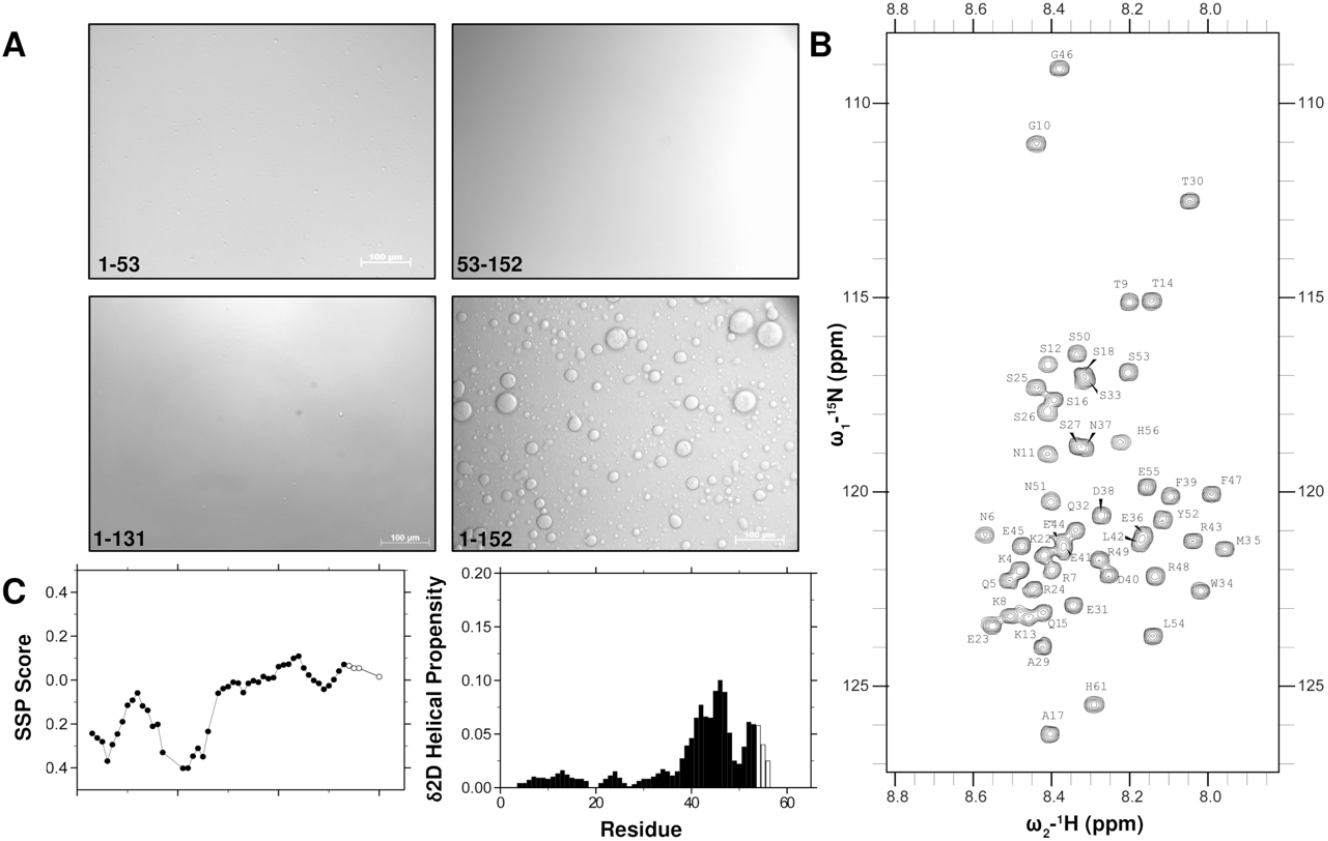
ORF1_1-152_ readily phase-separates, and the N-terminal domain and coiled-coil domain are both required for phase separation. **A.** Constructs of the N-terminal region (1–53), the coiled-coil domain (53–152), and truncation of the coiled-coil domain (1–131) in a tandem construct remain soluble at lower salt concentration (300 mM NaCl). In contrast, the full-length tandem construct lacking the RRM and CTD (1–152) phase separates to the same extent as the full-length protein, indicating that both the N-terminus and coiled-coil domains are necessary for phase separation. **B.** The narrow chemical shifts of ORF1_1-53_ in the ^15^N-^1^H HSQC experiment demonstrate that this region is intrinsically disordered. Resonances beyond 53 correspond to the histidine tag. **C.** The secondary structure propensity (SSP) score and δ2D Helical Propensity scores show that residues 41-49 have slight α-helical character. Open circles and open bars correspond to residues from the histidine tag.

**Table 1.** Summary of constructs used.

| ORF1 CONSTRUCT | DOMAIN ORGANIZATION | PHASE SEPARATION | RETROTRANSPOSITION |
| --- | --- | --- | --- |
| ORF1<sub>1-338</sub> | Full length | Normal | Normal |
| ORF1<sub>1-53</sub> | N-terminal region | None | N.D. |
| ORF1<sub>1-152</sub> | NT + coiled coil | Normal | N.D. |
| ORF1<sub>53-152</sub> | Coiled coil | None | N.D. |
| ORF1<sub>53-338</sub> | ΔNT | Weak | None (19) |
| ORF1<sub>66-338</sub> | ΔNT + truncated coiled coil | Weak | None (19) |
| ORF1<sub>157-252</sub> | RRM | None | N.D. |
| ORF1<sub>254-338</sub> | CTD | None | N.D. |
| ORF1<sub>157-338</sub> | RRM + CTD | None | N.D. |
| ORF1-338 K3A/K4A | FL | Normal | N.D.<br>ORF1-338 G2A/K3A/K4A: None (47)<br>ORF1-338 K3R: Increased (19) |
| ORF1-338 K3E/K4E | FL | Weak | N.D. |
| ORF1-338 K3A/K4A/R7A/K8A | FL | Weak | N.D. |
| ORF1-338 K3E/K4E/R7E/K8E | FL | Weak | N.D. |
| ORF1-338 R7E/K8E | FL | Weak | N.D. |
| ORF1-338 S27D | FL | Increased | Increased (41) |
| ORF1-338 L93P | FL | Increased | N.D.<br>ORF1-338 L93N/L199N: None (19)<br>ORF1-338 E89A/L90A/M91A: None (47)<br>ORF1-338 E92A/L93A/K94A: None (47) |

### The N-terminal 53 residues of ORF1 are intrinsically disordered but have helical propensity between residues 41 to 49

ORF1 residues 1 to 53 are predicted to be disordered using both long-range and short-range disorder prediction algorithms (IUPRED and SPOT-Disorder-Single (39, 40)), which is in agreement with the reported random coil signal of ORF1_1-51_ in circular dichroism experiments (19). To directly interrogate the structure of the ORF1 N-terminal region with residue-by-residue resolution, we used solution NMR spectroscopy on samples of ORF1_1-53_. The backbone amide resonances of ORF1_1-53_ are situated in a narrow chemical shift range of 8.6 to 7.9 ppm in ^1^H in the ^15^N-^1^H Heteronuclear Single Quantum Coherence (HSQC) correlation spectrum, which is characteristic of disordered proteins (Fig. 2B). The chemical shifts of Cα and Cβ are sensitive to partial secondary structure formation. Using two measures for predicting secondary structure from the observed chemical shift, we find slightly positive values in secondary shift perturbation (SSP) consistent with slight helical propensity (δ2D, ~10%) between residues E41 to R49 (Fig. 2C).

Because formation of partial structure should slow local reorientational motions in parts of ORF1, we acquired ^15^N spin relaxation data to examine the motion of ORF1_1-53_ at each residue position. The relaxation parameters are not uniform across the entire protein sequence and are significantly elevated after residue Q32 in two pH conditions. The average *R*_1_, *R*_2_, and heteronuclear NOE ratio parameters at pH 6.0 for residues of region 1-31 (1.32±0.12 s^−1^, 2.08±0.33 s^−1^ and −0.40±0.23) increase to average values of 1.74±0.15 s^−1^, 3.34±0.47 s^−1^ and −0.21±0.14 for residues 32-53, with a similar trend at pH 7.25. Together, these data are consistent with some partial structure for residues 32-53. We also acquired ^15^N-^1^H HSQC correlation spectra at varying ORF1_1-53_ concentrations. No chemical shift perturbations were observed with increasing protein concentrations from 20-700 μM, suggesting that the N-terminal region does not self-associate and exists as a monomer in solution (data not shown).

### The N-terminus of ORF1 is necessary, but not sufficient, for phase separation and interacts with the coiled-coil domain

Although deletion of the N-terminal region severely impairs LLPS, ORF1_1-53_ does not self-assemble in solution and is insufficient for phase separation alone. These observations imply that a component of the coiled-coil domain may be necessary to promote ORF1 condensation. Therefore, we used DIC microscopy to assay the ability of the individual domains to promote phase separation in *trans*. Titrating higher concentrations of the N-terminus into the coiled-coil domain did not promote phase separation up to a 1:6 molar ratio (Fig. 3A). However, titration of ORF1_1-53_ into full-length ORF1_1-338_ decreased phase separation in a concentration dependent manner, demonstrating that residues 1-53 can compete with the full-length protein for binding (Fig. 3B).

To address whether residue-specific interactions occur between the N-terminus and coiled-coil domain, we titrated ORF1_53-152_ into ^15^N isotopically enriched ORF1_1-53_. Addition of ORF1_53-152_ induces line broadening of the ORF1_1-53_ spectrum compared to free ORF1_1-53_. Peaks across the entire sequence, but especially in the transiently helical region following residue Q32 show significantly decreased intensities (Fig. 3C, D). Furthermore, the ^15^N *R*_2_ relaxation rates are elevated around residues M35, S50, and N51 (Fig. 3E, F). Because enhanced line widths and elevated transverse relaxation rates are consistent with either binding or conformational exchange, these data provide evidence that regions with lower signal intensity and higher relaxation rates correspond to residues contacting the coiled coil. It is important to note that the line broadening and enhanced ^15^N *R*_2_ are complementary and enhanced ^15^N *R*_2_ is not observed for every residue for which line broadening is observed. It is possible that regions from residues 32-53 contain the primary binding site while residues 1-31, which show reduced intensity but no large difference to ^15^N *R*_2_ relaxation, form transient contacts with the coiled-coil domain, leading to a chemical exchange induced line broadening. We hypothesized that residues 32-53 form the primary interaction sites that mediate initial binding with ORF1_53-152_; consequently, any changes within these sites should minimize the interaction between ORF1_1-53_ and ORF1_53-152_. The ^15^N-^1^H HSQC spectrum of a truncated ORF1_1-46_ construct is superimposable on the HSQC spectrum of ORF1_1-53_ in the unbound form, aside from the truncated region. At the same concentrations and conditions, significantly less line-width broadening is observed for ORF1_1-46_ than for ORF1_1-52_ in the presence of ORF1_53-152_, indicating that the interactions with the coiled-coil domain are partially disrupted by the removal of residues 47-53 (Fig. S2 in the Supporting Material). Based on these data, we hypothesized that ORF1 phase separation proceeds by transient, direct interactions between the N-terminal disordered region and the trimeric coiled-coil domain.

**Figure 3.**
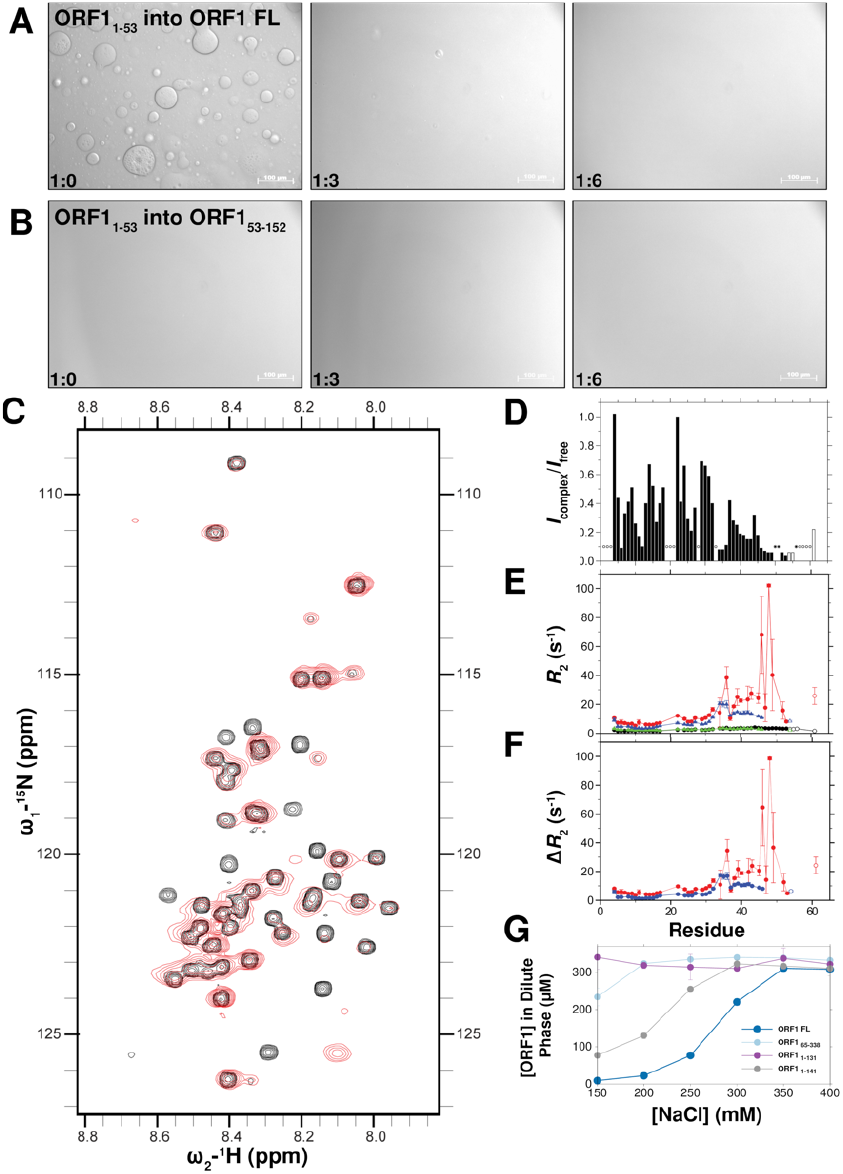
The ORF1 N-terminal domain interacts with the C-terminal 22 residues of the coiled-coil domain. A. ORF1_1-53_ can disrupt phase separation of the full-length ORF1_1-338_ protein in 300 mM NaCl, suggesting it competes for interaction sites found in ORF1_1-338_, but cannot promote phase separation of the coiled-coil domain in *trans*, as shown in **B**. This suggests that multivalency and interdomain interactions provided by the fusion of these domains is important for LLPS. **C.** Comparison of ^15^N-1H HSQC correlation spectra of 700 μM ORF1_1-53_ alone (black) and 700 μM ORF1_1-53_ mixed at an equimolar ratio with 700 μM ORF1_53-152_ (red) shows line broadening of ORF1_1-53_ in the presence of the coiled-coil domain with resonance attenuation consistent with binding of ORF1_53-152_ around residues 47-53 of ORF1_1-53_ and attenuations distributed across the entire sequence, shown in **D**, consistent with binding at multiple sites. Open bars correspond to residues from the histidine tag. Elevated values for the ^15^N *R*_2_ relaxation rate constants (**E**) are observed in the presence of ORF1_53-152_ at pH 7.25 (black) and pH 6.0 (red). Open circles correspond to residues from the histidine tag. **F**. The difference in *R*_2_ in the presence and absence of ORF1_53-152_, *ΔR*_2_, provides further evidence that residues 47-53 are perturbed by addition of ORF1_53-152_ due to slowed or conformational exchange because of binding to the structured, slow-moving coiled coil. **G**. Quantification of the ORF1 protein concentration remaining in the dispersed phase following centrifugation demonstrates that ORF1_1-131_ phase-separates much less readily than full-length ORF1_1-338_, while ORF1_1-141_ LLPS is intermediate. N-terminal domain deletion ORF1_65-338_ also significantly disrupts LLPS.

### Truncation of the C-terminal 22 residues of the coiled-coil domain abolishes phase separation

To further investigate the nature of the intermolecular interactions between the N-terminal disordered region and the coiled-coil domain, we asked if a head-to-tail orientation could facilitate the intermolecular contacts we observed by NMR spectroscopy. This type of interaction would place the N-terminal disordered region close to the base of the coiled-coil domain. We therefore generated two constructs that truncated the C-terminal 12 or 22 residues, ORF1_1-141_ and ORF1_1-131_, respectively. Gel filtration showed that both of these constructs still trimerize in solution. While ORF1_1-141_ showed decreased phase separation compared to ORF1_1-152_, ORF1_1-131_ showed no phase separation by DIC in the conditions and concentrations assayed, irrespective of salt concentration (Fig. 2A, Table 1). These results were further quantified by measuring the amount of protein remaining in the dilute phase in different buffer conditions compared to the full-length ORF1_1-338_ (Fig. 3G).

To address whether the residue-specific interactions observed between ORF1_1-53_ and the full-length coiled-coil domain ORF1_53-152_ were disrupted by removing the C-terminal 22 residues, we collected the ^15^N-^1^H HSQC spectrum of ^15^N isotopically enriched ORF1_1-53_ and added an equimolar ratio of ORF1_53-131_. In these conditions, we did not observe line width broadening (Fig. S3 in the Supporting Material) as we observed after addition of ORF1_53-152_. Thus, these data are consistent with a model where interactions between residues 46-53 and 131-152 are required for ORF1 phase separation.

### Mutations within both the N-terminus and coiled-coil domain modulate the properties of ORF1 phase separation

We next examined the contribution of specific residues or residue types to ORF1 phase separation. Previous mutagenesis and deletion studies showed that the basic residues at the extreme N-terminus are necessary for L1 retrotransposition in HeLa cells (14, 19, 41, 42) and that triple mutagenesis of G2, K3, and K4 to alanines or deletion of these residues inhibited retrotransposition to the same extent as the polymerase-dead mutant D702Y in ORF2 (19, 42). Unlike the disordered, low-complexity sequences observed in several phase-separating proteins, such as Fused in Sarcoma (FUS) and TDP-43, the ORF1 N-terminus is neither composed of repeat units nor highly enriched in a particular residue type (35, 43). Instead, ORF1 demonstrates charge patterning in the N-terminus with basic and acidic patches organized clusters throughout the sequence, akin to the RNA helicase Ddx4 (Fig. S4 in the Supporting Material) (44, 45). Due to our observation that increasing the ionic strength of the buffer outcompetes the protein-protein interactions leading to ORF1 phase separation, we hypothesized that weak, transient interactions at these positively and negatively charged blocks may contribute to ORF1 phase separation. Therefore, we investigated the effects of mutations within the N-terminal region that have been documented to either inhibit or enhance retrotransposition of the L1 element in context of full-length ORF1_1-338_. We generated double and quadruple mutants at positions K3, K4, R7, and K8 to assay whether changes in the charge patterning in the N-terminus affect ORF1 phase separation. Replacing lysine residues with glutamates in the K3E/K4E double mutant reduced the ability of ORF1 to phase separate in conditions that are favorable for the wild-type protein but did not completely abolish phase separation at lower salt concentrations (50-150 mM NaCl). In the quadruple mutants, removing the N-terminal basic patch (K3A/K4A/R7A/K8A) or introducing negative charges (K3E/K4E/R7E/K8E) also reduced ORF1 phase separation (Fig. 4).

**Figure 4.**
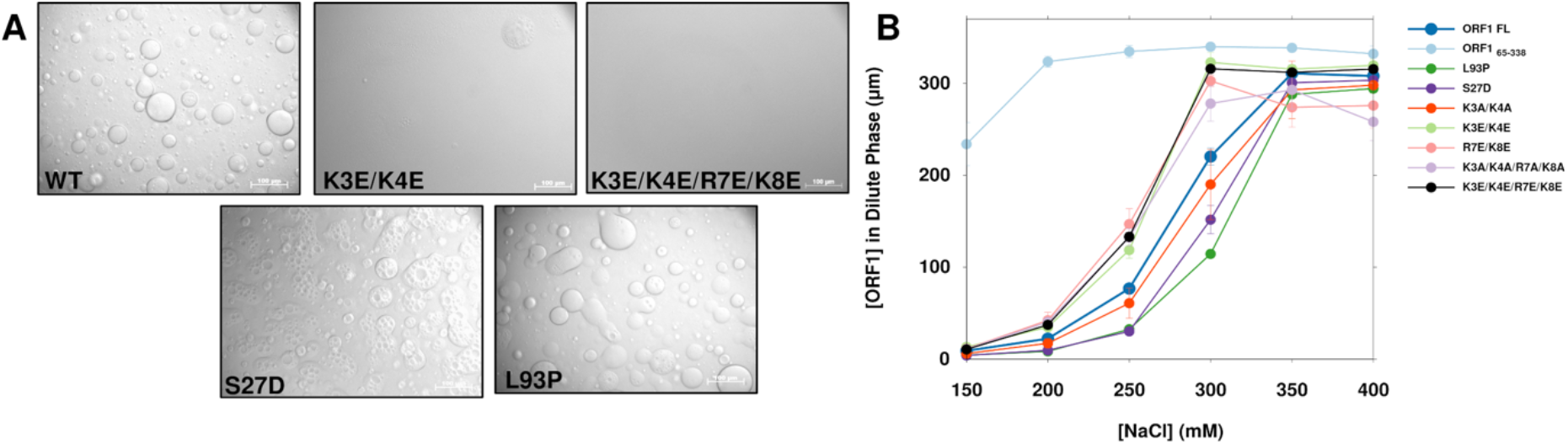
Mutations in both the N-terminus and coiled-coil domain modulate phase separation of the full-length protein. **A.** Altering the charge at the extreme N-terminus (K3E/K4E and K3E/K4E/R7E/K8E) impairs the ability of ORF1 to phase separate in 300 mM NaCl. Some mutations in the N-terminus and coiled-coil domain also appear to enhance the quantity and size of phase-separated droplets, as demonstrated by mutations S27D and L93P. **B.** Quantification of ORF1 protein concentration in the dilute phase shows that mutations that introduce a positive charge at the extreme N terminus significantly impair phase separation in comparison to the wild-type protein whereas the S27D and L93P mutations enhance phase separation.

In addition to the charge mutations, we examined the S27D phosphomimetic mutation because phosphorylation by proline-directed protein kinases increases L1 retrotransposition in cells (41). As observed by DIC microscopy, the S27D mutation increased the quantity and size of droplets compared to the wild-type protein, which is further substantiated by a reduced concentration of the S27D mutant protein in the dilute phase of the coexistence curve when compared to wild-type ORF1 (Fig. 4). Similarly, we tested a naturally occurring L93P mutation in the stammer of the coiled-coil domain (46) and found enhanced phase separation in comparison to the wild-type protein, demonstrating that mutations within the coiled-coil domain also modulate the behavior of ORF1 phase separation (Fig. 4). Taken together, these data suggest that mutations in both the charged residues and coiled-coil domain modulate liquidliquid phase separation.

## Discussion

### The disordered N-terminal domain promotes liquid-liquid phase separation

Prior to our observations of ORF1 phase separation, no function had been proposed for the ORF1 N-terminal region, despite the fact that a charged N-terminus is highly conserved across species and that extreme N-terminal residues are necessary for L1 retrotransposition (19, 47). Here, we demonstrate that the full-length ORF1 protein forms liquid droplets in a salt-dependent manner and that the N-terminal region is disordered but required for this phase separation because it forms contacts with the coiled-coil domain. Differences in ORF1 phase separation demonstrated by both salt dependence and mutational studies altering charged residues in the N-terminal region suggest that electrostatic interactions are important for driving coalescence into the phase separated state.

L1 is an ancient retroelement that has co-evolved with its eukaryotic hosts over vast stretches of evolutionary time. The ORF1 protein has adopted several functions and domains across evolutionary time in various species, including domain shuffling with functional host genes (48–50). Khazina and Weichenrieder defined two main classes of ORF1 proteins: the Type I family, found in plants, that contains an RRM and a CCHC zinc finger domain, and the Type II family found primarily in vertebrates, composed of a coiled-coil domain, RRM, and conserved CTD. In addition, an array of variations within each group has been documented that do not fit into these classifications (51). For example, the *Danio rerio* ORF1 protein has gained an esterase domain that may serve a membrane targeting function (52).

Human and mouse ORF1 belong to the Type II family and has been functionally and structurally well characterized. The RRM and CTD act together to bind RNA, and the coiled-coil domain functions to trimerize the protein. Both are required for biological activity, namely, to support the retrotransposition lifecycle of the L1 element. The function of the largely disordered N-terminal region (residues 1-53) has remained enigmatic, although mutagenesis studies have clearly shown it to be essential for viability. We propose that the biological function of the N-terminal region is to enable multivalent self-interactions that lead to *in vivo* particle assembly and LLPS. While the disordered N-terminal domain does not support LLPS alone, it acts in concert with the coiled-coil domain to enable LLPS of full-length ORF1. While our *in vitro* LLPS conditions are unlikely to fully recapitulate LLPS *in vivo*, by biophysically characterizing a number of deletions as well as single and multiple point mutations, we demonstrated a strong correlation between LLPS and biological function (retrotransposition).

### Sequence characteristics of the ORF1 N-terminal region

The N-terminal region shows the highest variability in length and sequence composition across species. However, a basic patch at the extreme N-terminus is highly conserved and is absolutely necessary for retrotransposition in human cells (19, 47). Though less conserved than the basic patch, the charge around residues 47-53, which interacts with the base of the coiled-coil domain, is well conserved across a number of vertebrates. Mutation of residues 44-46 (EEG) or 50-52 (SNY) to alanine reduces retrotransposition by approximately 50% compared to wild-type ORF1 in cells in both cases (47). Aside from the deletion or mutation of the first three residues (GKK, which reduces retrotransposition to 4.6%), these two mutations in the disordered N-terminus show the greatest reduction on retrotransposition (44-46: 58.5% and 50-52: 53.8%, (47).) This suggests that a reduction in the multivalent interactions seeded by the ORF1 N-terminus and coiled-coil domains may have a direct impact on the function of the L1 ribonucleoprotein particle.

The initial multivalency seeded by the interactions we observed using solution NMR spectroscopy between the base of the coiled-coil domains and the disordered N-terminus may be necessary to increase the local ORF1 concentration in order to nucleate phase separation. Once the concentration of ORF1 is sufficiently high, the intermolecular interactions mediated by the charged blocks within the ORF1 N-terminus then could establish a higher-ordered network that allows for demixing from the surrounding solution and provides environmental selectively for LLPS. The coiled-coil domain also displays charged residues on the solvent-exposed surface due to the organization of the R-h-x-x-h-E repeat motifs and the constellation of charged residues in the x positions (Fig. 1B) (19). While the arginine and glutamate at positions 2 and 6 of the repeat heptad are necessary for trimerization, the solvent-exposed charged residues may provide additional electrostatic interaction sites. The N-terminal domain may then form multiple electrostatic interactions with either the coiled-coil domain or possibly even N-terminal domains of neighboring ORF1 molecules in the phase separated state. Hence, our work shows that the charged region 47-53 contributes significantly to interactions with the coiled-coil domain leading up to phase separation but that charged residues near the N-terminus also play a role in stabilizing the phase-separated form.

K3R and L93P are two naturally occurring polymorphisms among the 146 intact and active human L1 sequences in the L1Base2 database (46). While these variants are competent for retrotransposition, removal of a single lysine at the N-terminus (K3) abolished retrotransposition in cell culture assays (19). The K3R polymorphism, in addition to maintaining the positive charge, may increase the interactions at the N-terminus due to the enhanced pi-character introduced with the arginine mutation and has been shown to increase retrotransposition of the L1 element to 157% compared to the wild type ORF1 protein (31, 53). The L93P polymorphism within the stammer (residues 91-93) may further destabilize the coiled-coil domain and allow for increased flexibility at the N-terminus of the coiled-coil domain itself which may help extend the disordered region when the beginning of the coiled coil is not fully populated. Khazina and Weichenrieder have demonstrated that an ORF1_1-103_ construct is unstructured by circular dichroism spectroscopy and that the L93N mutation, which is predicted to rigidify the coiled coil, decreases retrotransposition (19). Further, mutating the residues in the stammer to alanines, which would extend and stabilize the coiled coil, abolishes retrotransposition to the same extent as deleting or mutating the N-terminal basic patch (89-ELM-91: 4.18%, 92-ELK-94: 0.69%, (47).)

Alternating between the structured and unstructured states within the coiled-coil domain may provide a means for increasing the multivalency necessary for forming a phase-separated body. Elongating the disordered N-terminal region would allow for an extension of the N-terminus and increase any longer-range interactions with neighboring ORF1 molecules forming a higher-order mesh of the ORF1 protein. This would subsequently increase the local ORF1 concentration and allow for efficient bio-molecular condensation.

### ORF1 phase separation as a mechanism for RNA compaction

Phase separation of a number of RNA-binding proteins has been demonstrated throughout the literature, recently including efficient packing of virus particles. The measles virus utilizes phase separation to allow for rapid encapsulation and packing of the viral RNA (54). Additionally, the human immunodeficiency virus (HIV) nucleocapsid protein coordinates zinc into a phase-separated state which serves to increase viral replication (55). Even the SARS-CoV-2 nucleoprotein has been suggested to condense genomic RNA via phase separation (56). In a similar manner, the L1 RNP may utilize phase separation to package the L1 RNA, prevent host degradation of the transcript, and increase the replication potential of L1. While the L1 RNP does not package in a capsid, demixing from the surrounding cytosol may protect the L1 RNA intermediate from degradation by host factors, and compaction of the L1 transcript may enhance efficient nuclear transport of the assembled L1 RNP particle. Using RNA-fold, the predicted secondary structure of the L1 mRNA is extended with an enrichment of stem-loop structures at the ends of branch points, which would provide selective sites for ORF1 binding to the single stranded regions (57). Additionally, the ORF1 protein has been shown to form dumbbell shaped structures by atomic force microscopy (AFM) with an average length of 320 Å connecting the wider, distinct C-termini of the ORF1 trimers (58). It is plausible that ORF1 is able to bind the L1 RNA within single stranded regions and, through head-to-tail interactions between the N-terminus and the coiled-coil domains, as we observed using NMR spectroscopy, condense the transcript and sequester it into a coacervated state that encompasses the full L1 RNP assembly.

## Conclusion

Our data provide insights into the phase separation propensity and multivalent interactions between the N terminus and coiled-coil domain and suggest a potential role for the structural malleability encoded within the stammer region of the coiled-coil domain. Further experiments will need to explore phase separation of the L1 RNP in cells and whether disrupting ORF1 phase separation directly impairs retrotransposition of the L1 element *in vivo*. Pharmacological inhibition phase separation has emerged as a potential therapeutic approach (59) and modulation of ORF1 phase separation could provide an exciting novel strategy to interfere with L1 retrotransposition in aging and disease.

## Author Contributions

J.M.S. and G.J. conceived the study. J.C.N., M.T.N, N.L.F, J.M.S. and G.J. designed the experiments. J.C.N., G.Y.L. and M.T.N. performed the experiments. All authors contributed to data analysis and interpretation. J.C.N. wrote the manuscript with input from all authors.

## Acknowledgements

J.M.S. is a cofounder of Transposon Therapeutics, Inc., serves as Chair of its Scientific Advisory Board, and consults for Astellas Innovation Management LLC, Atropos Therapeutics, Inc. and Gilead Sciences, Inc. N.L.F. consults for and serves on the Scientific Advisory Board of Dewpoint Therapeutics.

