## Supporting Materials for "Phase separation of the LINE-1 ORF1 protein is mediated by the N-terminus and coiled-coil domain"

**E. coli codon optimized ORF1 gene sequence used in this study**

ATGGGTAAAAAACAGAATCGTAAGACCGGTAACAGCAAAA
CCCAAAGCGCGAGCCCGCCGCCGAAGGAACGCAGCAGCAGCCCGG
CGACCGAGCAGAGCTGGATGGAAAACGACTTCGATGAGCTGCGTG
AGGAAGGTTTTCGTCGTAGCAACTACAGCGAGCTGCGTGAAGACA
TCCAAACCAAGGGCAAAGAGGTGGAAAACTTTGAAAAGAACCTGG
AGGAATGCATCACCCGTATTACCAACACCGAGAAGTGCCTGAAAG
AGCTGATGGAACTGAAGACCAAAGCGCGTGAACTGCGTGAGGAAT
GCCGTAGCCTGCGTAGCCGTTGCGACCAGCTGGAGGAACGTGTGA
GCGCGATGGAGGATGAAATGAACGAGATGAAGCGTGAGGGTAAAT
TCCGTGAGAAGCGTATCAAACGTAACGAACAGAGCCTGCAAGAGA
TTTGGGATTACGTTAAGCGTCCGAACCTGCGTCTGATCGGTGTGC
CGGAGAGCGACGTTGAAAACGGCACCAAACTGGAAAACACCCTGC
AGGATATCATTCAAGAGAACTTTCCGAACCTGGCGCGTCAAGCGA
ACGTGCAGATCCAAGAAATTCAGCGTACCCCGCAACGTTATAGCA
GCCGTCGTGCGACCCCGCGTCACATCATTGTGCGTTTCACCAAGG
TTGAGATGAAGGAAAAAATGCTGCGTGCGGCGCGTGAGAAAGGTC
GTGTTACCCTGAAGGGCAAACCGATTCGTCTGACCGCGGATCTGA
GCGCGGAAACCCTGCAGGCGCGTCGTGAGTGGGGTCCGATCTTCA
ACATTCTGAAGGAGAAGAACTTTCAACCGCGTATCAGCTACCCGG
CGAAACTGAGCTTCATTAGCGAGGGCGAAATCAAGTACTTCATCG
ACAAGCAGATGCTGCGTGATTTCGTTACCACCCGTCCGGCGCTGA
AGGAGCTGCTGAAAGAAGCGCTGAATATGGAACGCAATAACCGCT
ACCAACCGCTGCAAAATCACGCGAAAATGTAA

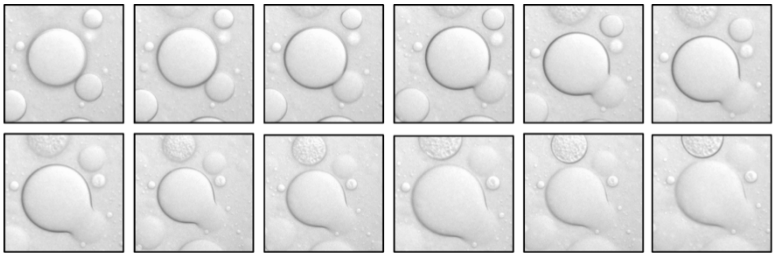

**Figure S1. ORF1 droplets behave as liquids.**
Images were taken in succession (within approximately 10 seconds) to demonstrate that the ORF1 droplets are capable of flowing and fusing in solution.

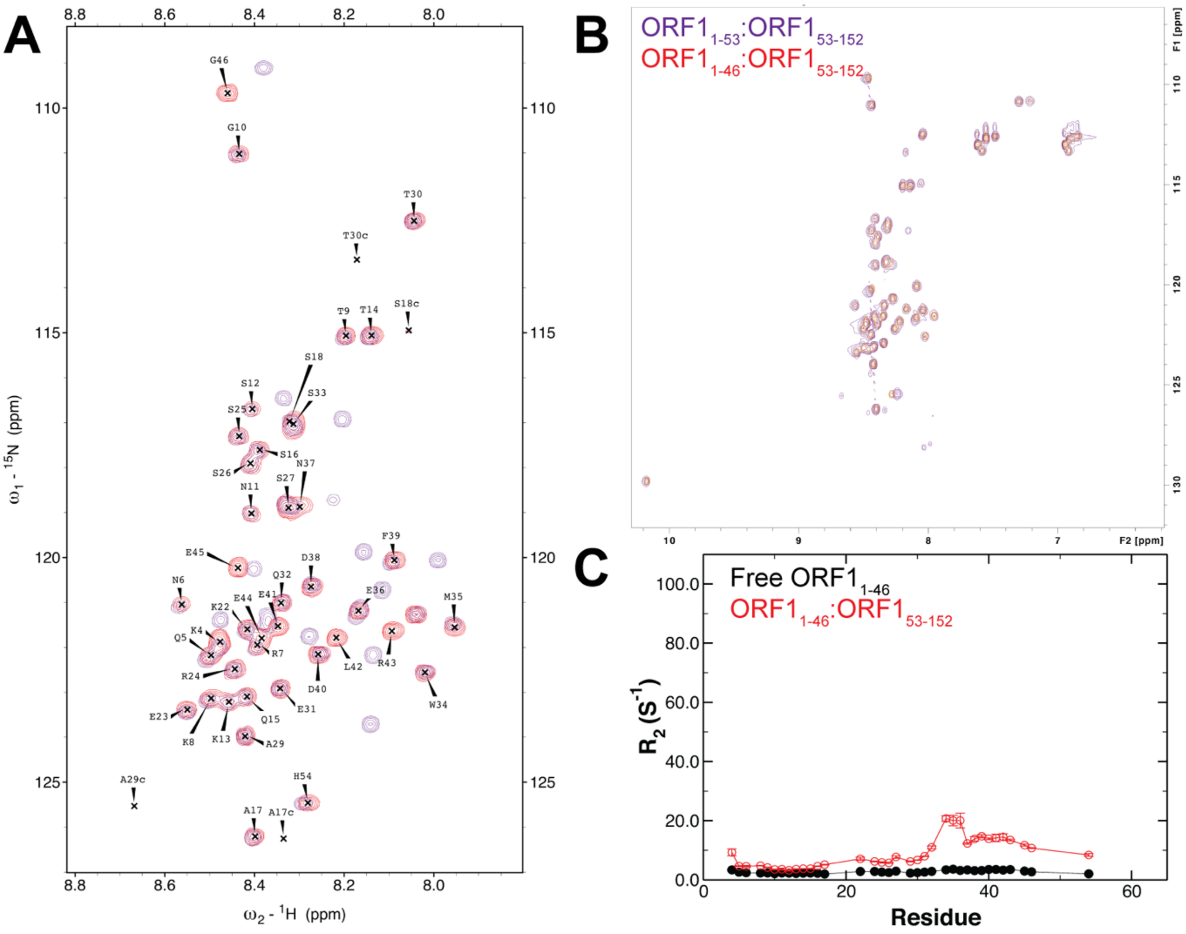


**Figure S2. ORF1$_{1-46}$ remains disordered but cannot interact with ORF1$_{53-152}$.**
**A.** The $^{15}$N-$^{1}$H HSQC correlation spectra overlay nicely between ORF1$_{1-53}$ (purple) and ORF1$_{1-46}$ (red). **B.** ORF1$_{1-46}$ no longer displays significant line broadening in the presence of ORF1$_{53-152}$ (red) as observed with ORF1$_{1-53}$ (purple), indicating that the interaction is disrupted by removing residues 47-53. **C.** The values of $R_2$ are slightly elevated in ORF1$_{1-46}$ in the presence of ORF1$_{53-152}$ (red) compared to free ORF1$_{1-46}$ (black), unlike the highly elevated values observed for ORF1$_{1-53}$ in the presence of ORF1$_{53-152}$. Open circles correspond to residues from the histidine tag. These data were acquired at 700 μM ORF1$_{1-53}$ or ORF1$_{1-46}$ and their respective 1:1 molar ratio complexes with 700 μM ORF1$_{53-152}$ (red).

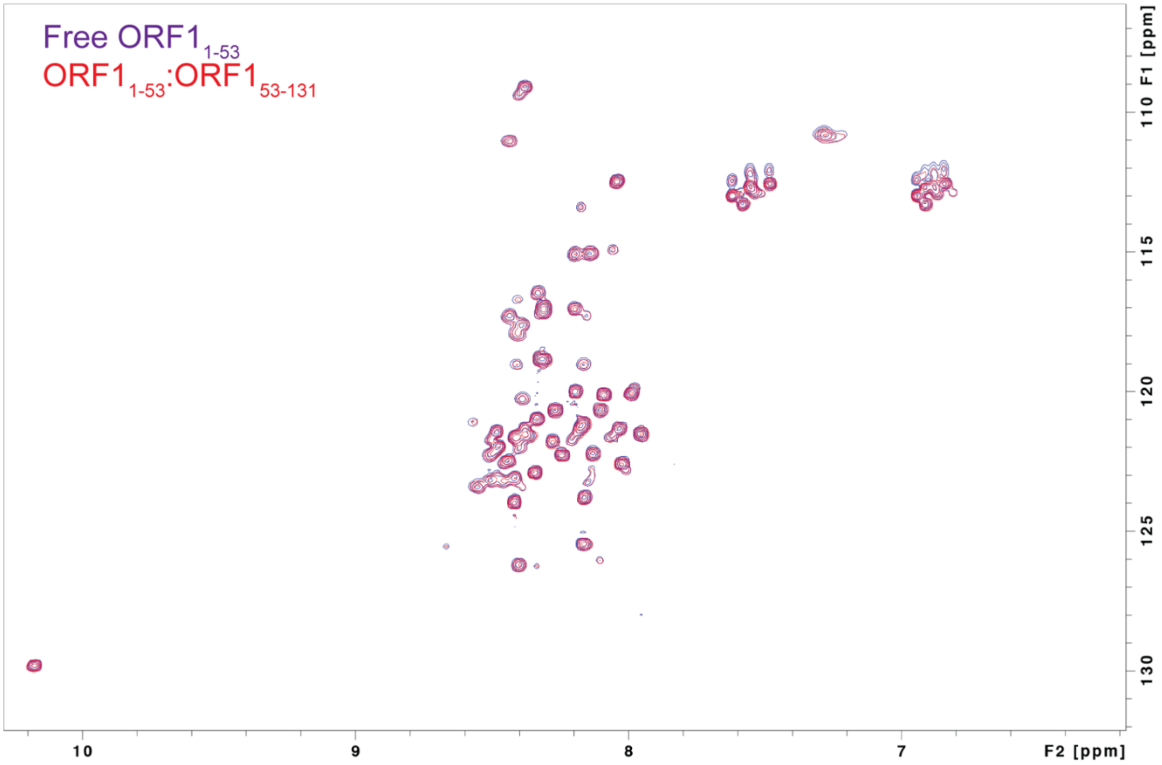


**Figure S3. ORF1$_{1\text{-}53}$ cannot bind to the C-terminal portion of the coiled-coil domain after truncating residues 132-152.**
No differences are observed between the $^{15}N$-$^{1}H$ HSQC correlation spectra of free ORF1$_{1\text{-}53}$ (790 μM) and ORF1$_{1\text{-}53}$ (300 μM) in the presence of the ORF1$_{53\text{-}131}$ (300 μM) truncated coiled-coil domain, demonstrating that the C-terminal 22 residues are necessary for that interaction.

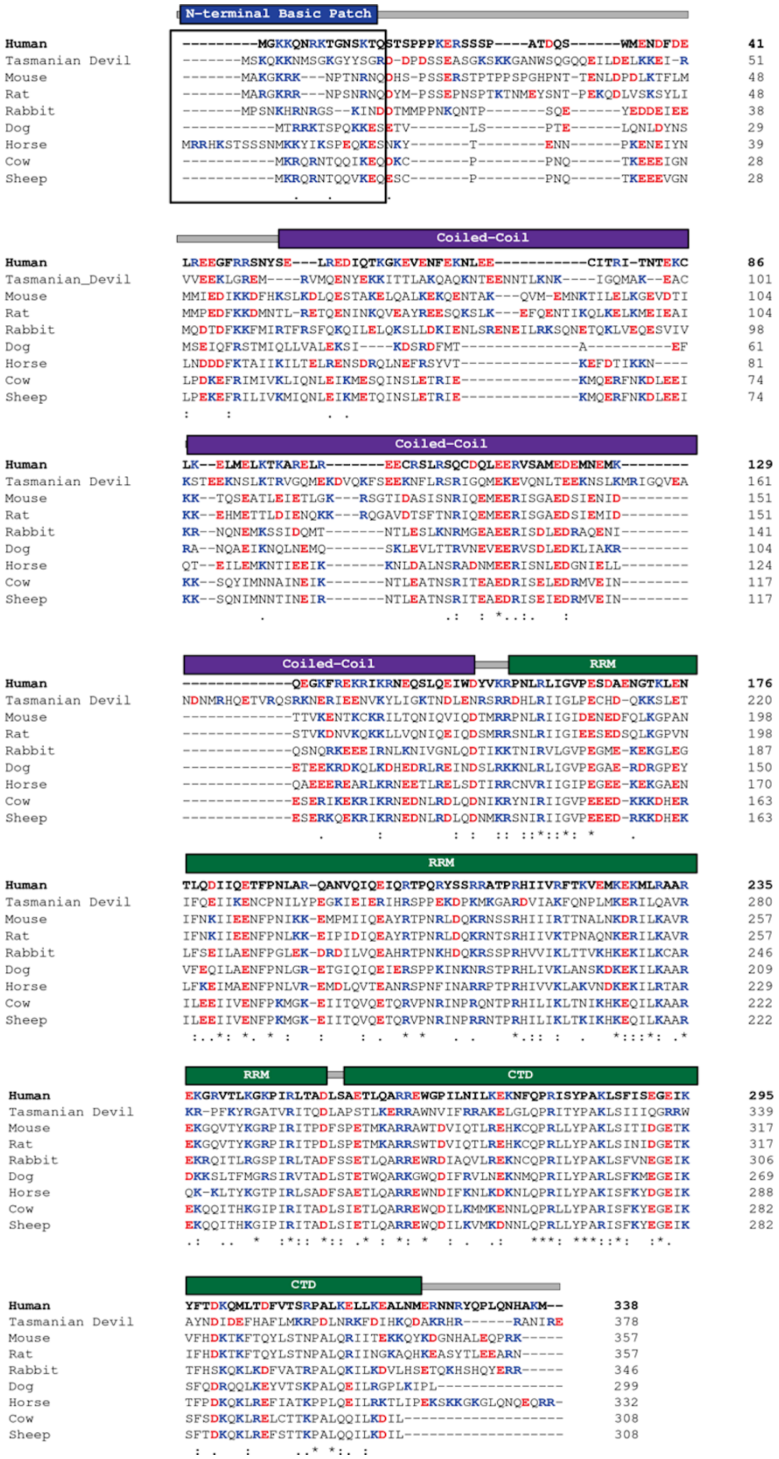


**Figure S4. The basic residues at the N-terminus of ORF1 are highly conserved.**
Accession numbers for the sequences are as follows: Human, AAA36590.1; Mouse, P11260.2; Tasmanian Devil, NW 003849619.1; Rat, AAY88219.1. The remaining sequences were obtained from L1Base2 (1). Domain architecture placed above the alignments corresponds to the human ORF1 do main boundaries.
